# A Novel Mouse Wound Model for Scar Tissue Formation in Abdominal-Muscle Wall

**DOI:** 10.1101/2021.08.05.455355

**Authors:** Shiro Jimi, Arman Saparov, Seiko Koizumi, Motoyasu Miyazaki, Satoshi Takagi

**Affiliations:** Central Lab for Pathology and Morphology, Faculty of Medicine, Fukuoka University, Fukuoka 814-0180, Japan; Department of Medicine, School of Medicine, Nazarbayev University, Nur-Sultan 010000, Kazakhstan; Nitta Gelatin Inc. R&D Center, Osaka 581-0024, Japan; Department of Pharmacy, Fukuoka University Chikushi Hospital, Fukuoka 818-0067, Japan; Department of Plastic Reconstructive and aesthetic Surgery, Faculty of Medicine, Fukuoka University, Fukuoka 814-0180 Japan

**Keywords:** Animal model, Fibrosis, Granulation tissue, Hypertrophic scar, Scaring, Wound healing

## Abstract

Scar tissue formation is a result of excess healing reactions after wounding. Hypertrophic scars scarcely develop in a mouse. In the present study, we established a novel experimental model of a scar-forming wound by resecting a small portion of the abdominal wall on the lower center of the abdomen, which exposed contractive forces by the surrounding muscle tissue. As a tension-less control, a back-skin excision model was used with a splint fixed onto the excised skin edge, and granulation tissue formed on the muscle facia supported by the back skeleton. One week after the resection, initial healing reactions such as fibroblast proliferation took place in both models.

However, after 21 days, lesions with collagen-rich granulation tissues forming multiple nodular/spherical-like structures developed only in the abdominal-wall model. The lesions are analogous to scar lesions in humans. Such lesions, however, did not develop in the back-skin excision model. Therefore, this animal model is unique in that fibrous scar tissues form under a physiological condition without using any artificial factors and is valuable for studying the pathogenesis and preclinical treatment of scar lesions.

**Summary Statement:** Scar lesions are hardly developed in animals. We thus developed a scar lesion in mice without using any artificial factors. The model is reliable, reproducible, and valuable.

## Introduction

The best outcome of wound healing is that the damaged tissue foci substitute with original tissue structures. The wound healing process is divided into four stages (Rodrigues et al., 2019; Zomer and Trentin, 2018) including coagulation, inflammation, proliferation, and maturation. Proper healing can be accomplished only by efficiently passing through all of these stages in order. In particular, granulation tissue formation and regeneration are fundamental cellular events in the proliferation stage. Different types of cells participate in this stage (Greenhalgh et al., 1990); fibroblasts produce extracellular matrix (Tracy et al., 2016), myofibroblasts enable wound contraction (Vallee and Lecarpentier, 2019), and endothelial cells establish a vascular network (Velnar and Gradisnik, 2018). Self-forming collagen-rich scaffolds support the acceleration of regenerative processes, in which granulation tissue finally regresses after the end of wound healing. However, a decrease in granulation tissue’s quantity and quality by malnutrition and defect of neovascularization can cause severe problems in wound healing, resulting in intractable wounds. Moreover, disorders that affect regeneration can cause a decrease in organ functions and appearance.

Collagen is an extracellular protein and plays a significant role in providing physical tissue strength and maintenance (Tracy et al., 2016). Collagens produced by fibroblasts play a cellular scaffold in granulation tissue. In the matrix, endothelial cells donate neovascularization, and keratinocytes induce epithelial healing. Granulation tissue develops after tissue damage not only in the skin but also in other organs. If the granulation tissue becomes fibrous with collagen accumulation during wound healing, scarring may appear with collagen hyalinization by unknown mechanisms. Responsible factors for the genesis of scar lesions are growth factors including transforming growth factor-β (TGF-β) (Liarte et al., 2020; Vallee and Lecarpentier, 2019), tensile forces, and intracellular SMAD-pathway activation via mechanoreceptor involving mechanisms (Harn et al., 2019). In the skin, an elevated lesion accompanied by massive fibrosis is called a hypertrophic scar (Ogawa, 2017), and it forms slowly after wounding in the skin turgor around the joints. The genesis of scar lesions is thus supposed to be phenotypic modulation of scar-forming cells under a hyper tensile force and delayed healing due to chronic inflammation.

Keloid also appears as an elevated lesion (Limandjaja et al., 2020); however, its lesion invades over the adjacent normal skin due to a higher proliferation potency. Like the hypertrophic scar, keloid occurs in the skin areas under a higher tensile force (Ogawa, 2017), yet, there is also a predisposition in certain races and patients. The emergence of proliferation-prone or variant cells is essential for keloid genesis, although its pathogenesis and factor(s) that are involved in this process are not fully deciphered.

Wound healing studies using animals have been carried out for many decades (Ahn and Mustoe, 1990; Falanga et al., 2004; Reid et al., 2004). Abercrombie et al. (1960) (Abercrombie et al., 1960) advocated that the skin with wounds in animals should be splinted to closely resemble human wound healing. Other investigators also indicated that skin mobility in animals affects wound contraction (Carlson et al., 2003; Davidson et al., 2013; Galiano et al., 2004; Kennedy and Cliff, 1979; Wang et al., 2013). Therefore, the applicability of animal wounds is an essential concern to human wound healing in basic science.

Animal models are crucial for the preclinical examination of human diseases. Animal wound healing models have been used to explore the pathogenesis and effectiveness of treatments in vivo under physiological conditions. Scar lesions after skin excision are nevertheless challenging to produce in animals, especially in rodents due to the looseness and tensile-less nature of the tissue. On the other hand, researchers have tried to generate animal models with hypertrophic scars using mice, pigs, and other animals (Ahn and Mustoe, 1990; Blackstone et al., 2017; Momtazi et al., 2013; Zhou et al., 2019). In 2007, Aarabi et al.(Aarabi et al., 2007) established a hypertrophic scar-forming mouse model by using a biomechanical loading device. Marchesini et al. (2020) (Marchesini et al., 2020) created a foreign body reaction-associated fibromuscular granulation tissue in rats. Liu et al. (2017) (Liu et al., 2017) created a pulmonary fibrosis model by using bleomycin in mice. However, no model for scaring after wounding in rodents has not been developed under a physiological condition without using any devices or chemicals.

We have therefore developed a scar-forming mouse model by resecting a part of the abdominal wall of muscle layers without using any devices, resulting in the development of granulation tissue constantly loaded by the surrounding muscle tension. As a result, a unique fibrous scar lesion subsequently developed. We also compared it to the tension-less granulation tissue formed on the back skeleton using the back-skin excision model (Jimi et al., 2020a; Jimi et al., 2017b; Jimi et al., 2017c; Jimi et al., 2020b).

## Results

### Wound contraction and tensile force

Two different wound models were utilized, which include the back-skin splint model (Figure 1A) where granulation tissue formed after skin excision on the back muscle facia supported by the back skeleton. The tension on the developing granulation tissue was minimized by not only the back skeleton, but also the splint (blue dot-line) fixed under the edge of the skin. The other model was the abdominal-wall excision model (Figure 1B). The abdominal muscle wall has no skeletal support. A small portion of the abdominal muscle wall was resected and covered by the skin. After wounding, no difference was found in body weight, WBC, and RBC between the groups during the study (Table S1), and wounds visually contracted in both the back-skin model (Figure 1A) and in the abdominal wall model (Figure 1B). On day 7, the outer look of the wounds on the abdominal wall showed a vertical oval shape from the abdominal cavity mucosa, but apparent raw lesions were scarcely visible after 21 days of the study.

**Figure 1.**
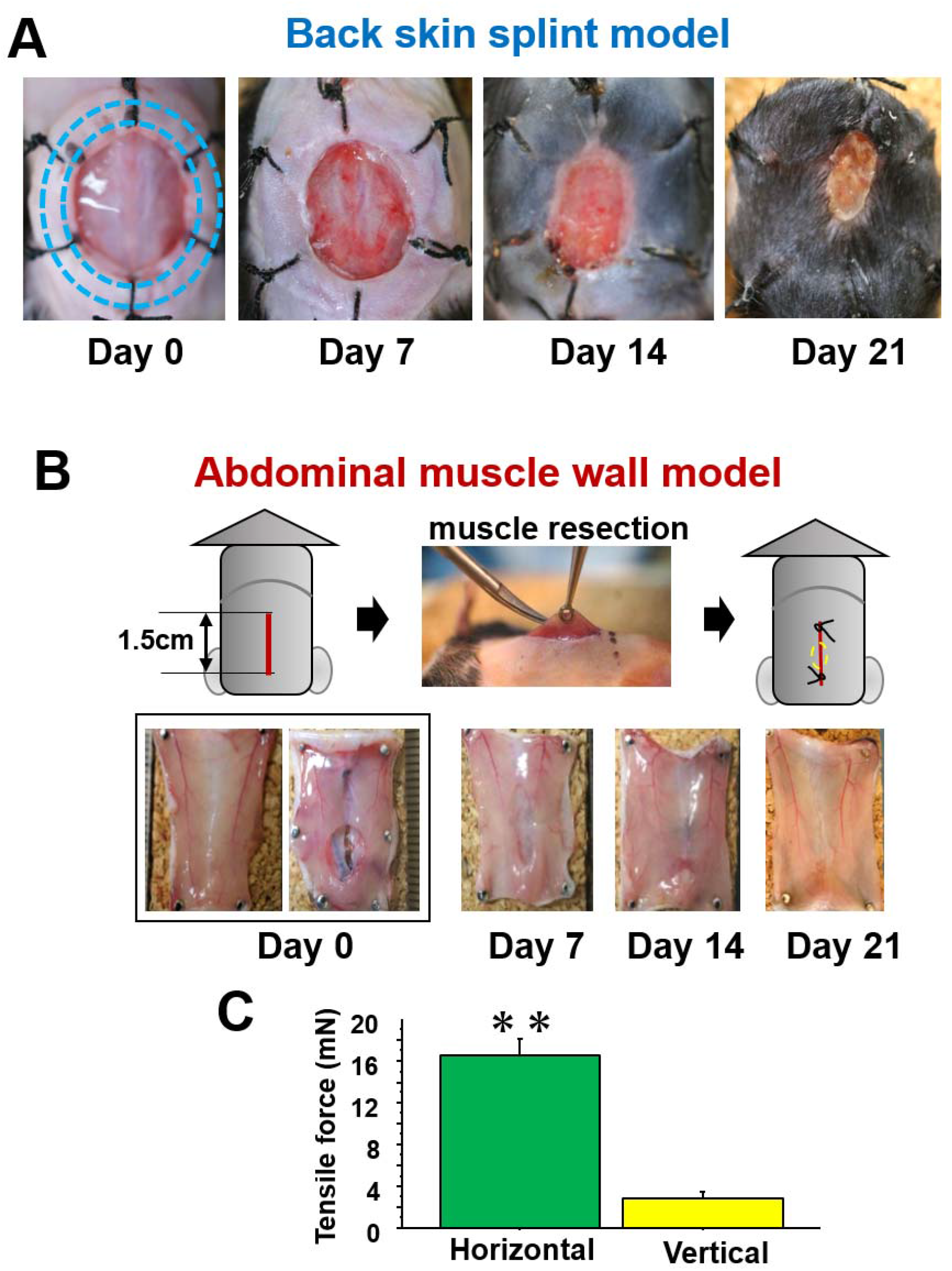
Two different models for wound healing with and without tensile force. A: The outer wound appearances by time in the back-skin splint model with tensile force-unloaded granulation tissue. Skin excised is fixed with the splint (blue dot-line) under the dermis, and granulation tissue forms on the muscle facia supported by the back skeleton. B: The inner wound appearance by time in the abdominal muscle wall model with tensile force-loaded granulation tissue. C: To analyze the tensile force loaded on the abdominal muscle wall, the wall was excised horizontally or vertically along a one-cm straight line, and tensile force was measured by the method described in the Materials and Methods section.

The tensile force in the abdominal wall without the skin was analyzed as a static physical force in the mice under general anesthesia. The abdominal muscle wall was resected 1 cm in a horizontal or vertical line. The vertical cut revealed a slight tensile force, and more than four-times greater tensile force was in the horizontal cut than the vertical cut (P<0.01) (Figure 1C). On the other hand, non of the tensile force was detected in the abdominal skin that was similarly cut in size and direction (data not shown).

### Development of granulation tissues

Granulation tissues formed on the wound in both models after wounding. In the back-skin model, the granulation tissue started under the edge of the skin excised and formed on the facia (Figure 2A, MT stain). On the other hand, in the abdominal model, the granulation tissue started to form from the excised abdominal muscle wall. On day 7, fibroblasts arranged parallel to the wounds (Figure 2B), showing a zonal structure that is similar in both models. In the back-skin model, the zonal growth pattern was maintained during the study (Figure 2A), except for an increase in collagen matrix accumulation. On the other hand, in the abdominal model, granulation tissue was arranged in a rippling pattern on day 14 (Figure 2B, middle). The lesions changed in a fibrous nodular/spherical structure on day 21 (Figure 2B, right). The area of collagenous matrix stained by MT stain increased with time in both models (Figure 2C), and its extent in the abdominal model was about 10-times greater than the back-skin model. We also checked the distribution of CD31-positive endothelial cells and α-SMA-expressing cells including myofibroblast in the well developing granulation tissues in both models on day 14 (Figure 2D). As compared to the back-skin model, CD31-positive neovessels were distributed to a lesser extent in the abdominal granulation tissue. However, α-SMA expressing cells were densely distributed along with the wavy extracellular matrix in nodular granulation tissue in the abdominal model.

**Figure 2.**
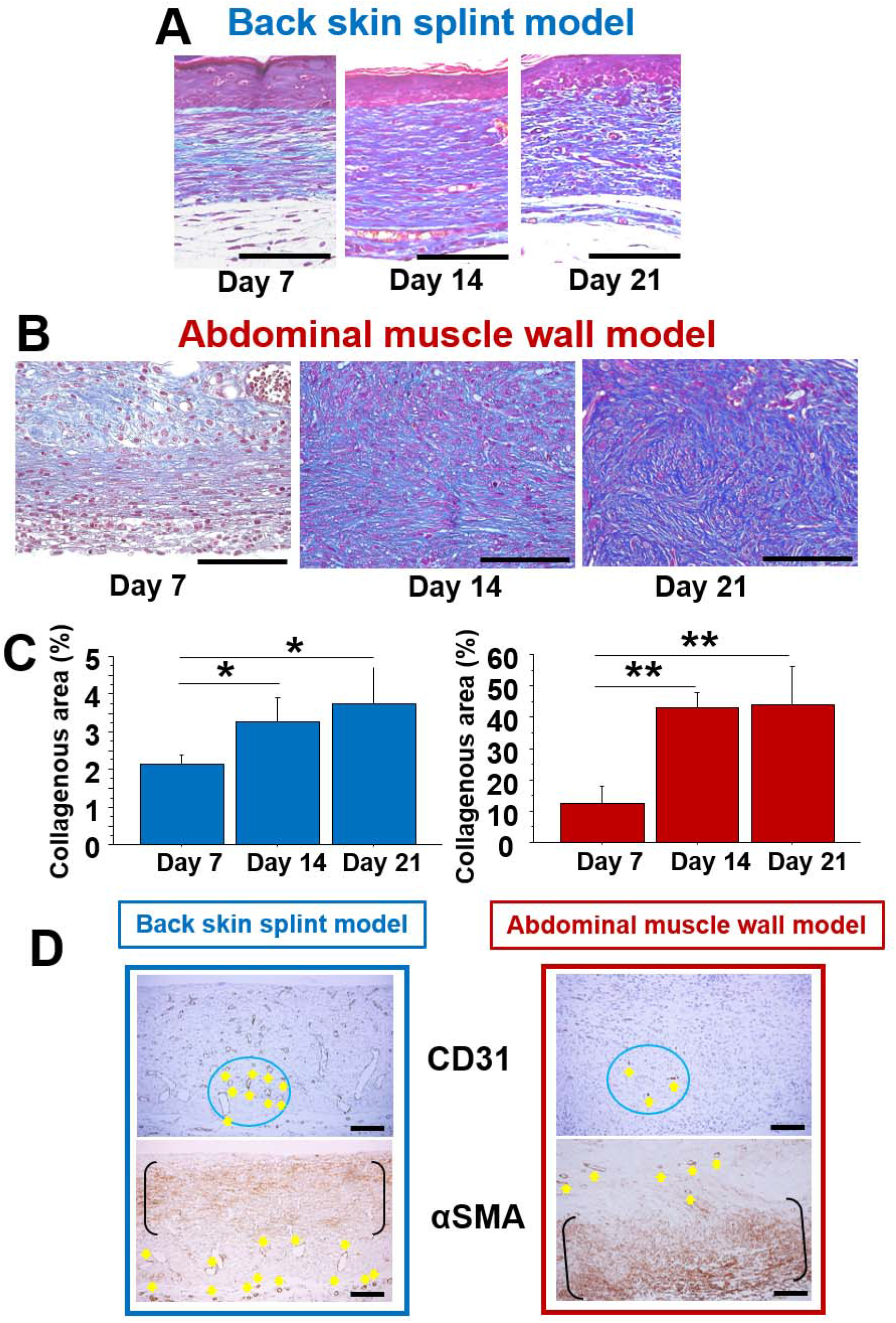
Cross-sections of the granulation tissue formed in the two different wound models and the histological characteristics. A: The granulation tissues were developed for 21 days in the back-skin splint model (MT stain). The pictures of the representative granulation tissues were taken under the active part of the epithelial regeneration. The granulation tissues maintained a zonal structure with parallelized fibroblasts during the study. Bars = 100 μm. B: Representative granulation tissues developed for 21 days in the abdominal-wall excision model (MT stain). The granulation tissue changed from a zonal structure (day 7) to a wavy nodular/spherical structure (days 14 and 21). Bars = 100 μ C: The comparison of the collagenous area in the developing granulation tissues on day 14 in the back-skin model (left) and the abdominal wall model (right). D: The representative IHC pictures for CD31 and α-SMA in the granulation tissue on day 14 from two different models. CD31-positive vessels in the circle area are shown by the arrows. α-SMA-positive cells distribute zonally between the parenthesis, and vessel walls are also positive for α-SMA (arrows). Bars = 100 μm.

### Excised tissue weight and wound area

After excising a part of the abdominal muscle wall in the abdominal model (Figure 3A), tissues weight and wound area were measured. The distribution of wound areas and tissue weights was 4.8∼25.1 mm^2^ and 2.2∼7.8 mg, respectively (Figure 3B). There was a significant correlation between these parameters (p<0.0001) (Figures 3C and 3D). Thus, the tissue weight could be used as a value for the extent of wounding damage.

**Figure 3.**
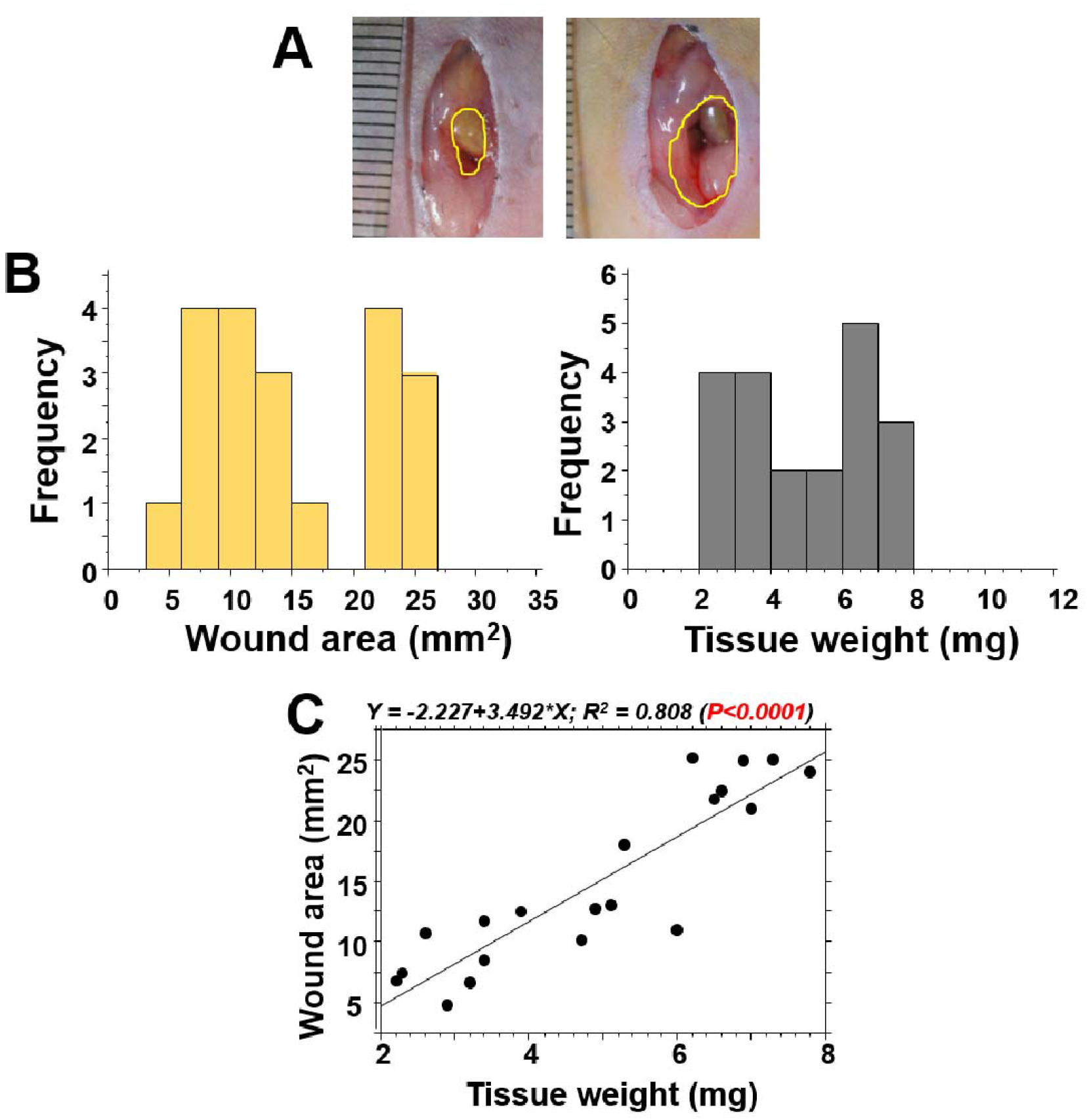
Wound area and tissue weight after resecting the abdominal wall. In 20 mice, a part of the abdominal wall was resected on day 0, as the abdominal wall excision model. A: The small and large areas of excised wounds: about 5 mm^2^ and 25 mm^2^, respectively. B: The frequency distributions of the wound area (left) and tissue weight (right). C: At the time of wounding, obtained tissue weight significantly correlated with the area of penetrated wound hole.

### Morphometrical analysis of granulation tissue

Figure 4A shows a representative wound picture of the abdominal wall on day 21 in the mouse with abdominal muscle resection and is a cross-sectional image. A granulation tissue developed between the abdominal muscle wall and wound surface exposed to the mucosa of the abdominal cavity. The granulation tissue shown by a yellow line and lateral wound length shown by a green line were morphometrically measured as granulation tissue area and granulation tissue length, respectively. Moreover, using Mt stain, the collagenous area stained by blue and the cellular area stained by red were also morphometrically measured (Figure 4B).

**Figure 4.**
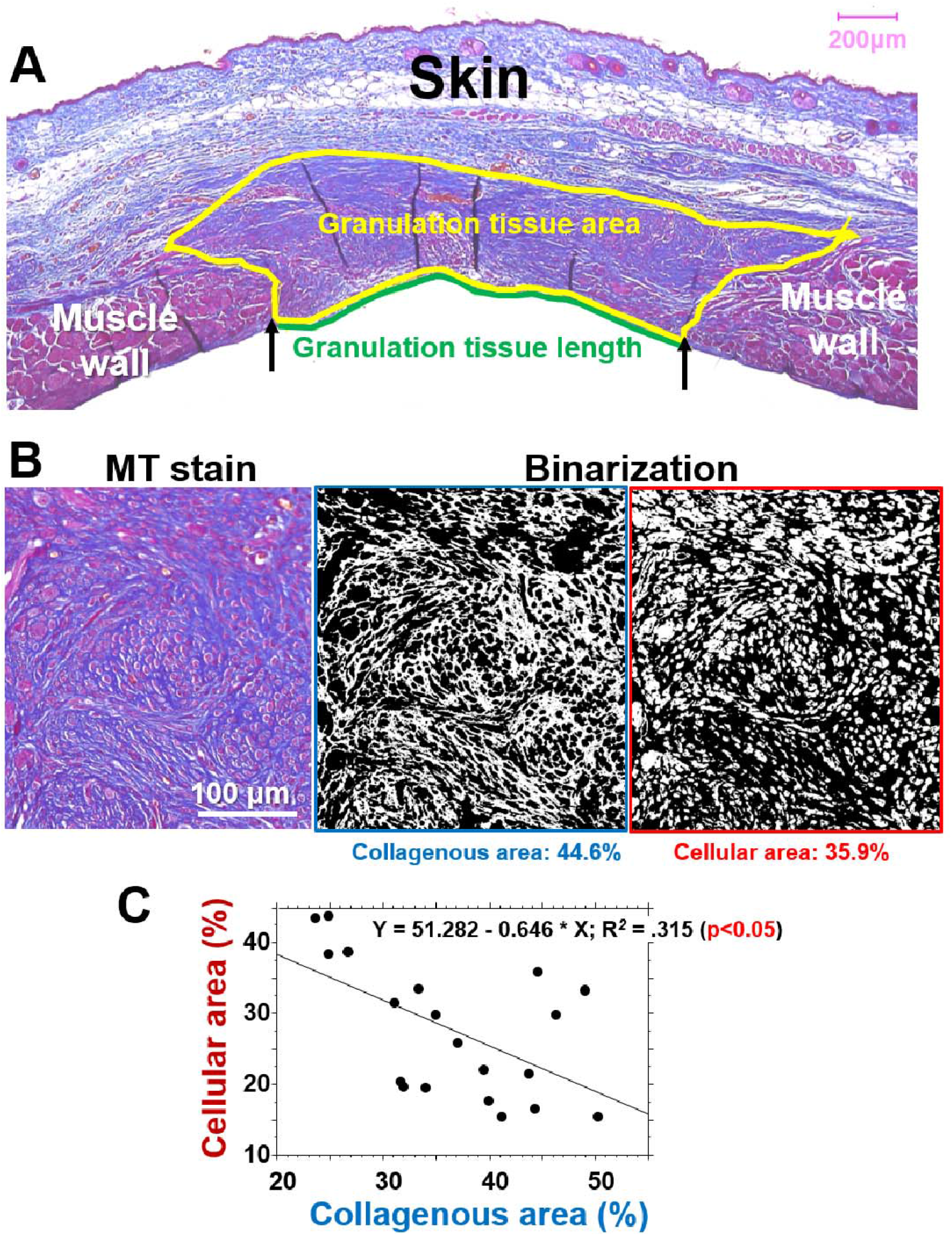
The histological evaluation of granulation tissue, including collagen and cellular elements. A: In the case of an abdominal wound in a cross-section on day 21 stained by MT stain: the granulation tissue developed between the abdominal muscle wall was estimated morphometrically by its area (yellow line) and the surficial length of the abdominal cavity (green line). B: Representative granulation tissue stained by MT stain, in which collagenous matrix is stained in blue, and the cells are stained by red. These elements were binarized and morphometrically measured. C: In the abdominal wall model, a negative correlation was found between the collagenous matrix area (%) and cellular area (%) in the granulation tissue on day 21.

### Development of granulation tissues with fibrosis

Collagenous and cellular areas stained blue and red in the MT stain, respectively, were quantified in the granulation tissue on day 21 (Figure 4B). The collagenous area was conversely correlated with the cellular area in the granulation tissue (p<0.05) (Figure 4C, left). It showed that fibrosis in the developed granulation tissue progressed following the decrease in cellularity.

### Histological characteristics of the granulation tissue and scar tissue

After the end of the study on day 21, different wound lesions developed (Figure 5). The granulation tissues developed were composed of the cellular elements, including fibroblasts, endothelial cells, adipocytes, and regenerative muscle cells, and extracellular matrix, including collagens. Using five values of tissue index, including tissue weight (TW) and wound area (WA), granulation tissue area (GA), granulation tissue length (GL), and collagen area (CoA), representative images of different sized granulation tissues are shown in Figure 5. These results indicated that collagen-rich fibrosis (scaring) advanced according to the increase in lesion size.

**Figure 5.**
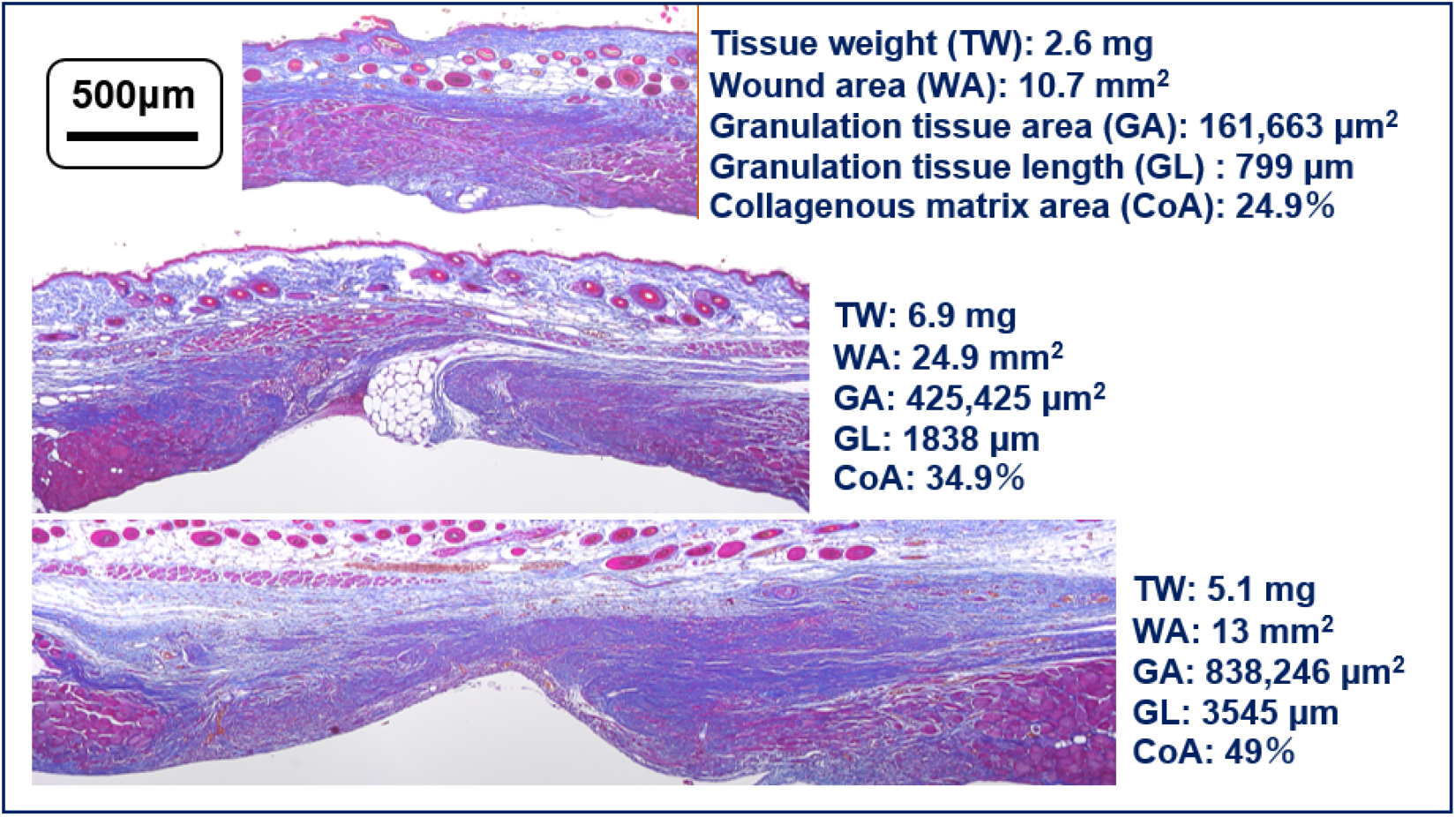
Differently structured lesions on day 21 in the abdominal wall model. Among the mice in the abdominal wall model, differently structured lesions developed. Obtained tissue indexes in each lesion are shown, including tissue weight (TW), wound area (WA), granulation tissue area (GA), granulation tissue length (GL), and collagenous matrix area (CoA). Collagen-rich fibrosis (scaring) advanced according to the increase in lesion size.

Nodular/spherical structures in the granulation tissue were formed (Figure 6A), and such lesions developed up to about 80% of mice. In advanced fibrous lesions with less cellularity, collagen matrices were hyalinized, showing as eosinophilic-hazy staining (Figure 6B). Collagens in the granulation tissue were further examined using the samples stained by picrosirius red stain and observed under polarization microscopy: The fibrous lesion developed in the abdominal wall model forms a storiform pattern (Figure 6C, left), and thickened collagen bundles in orange color (type | collagen) primarily observed, which differed from the thin collagens in green color (type ||| collagen) that accumulated in the granulation tissue in the back-skin model (Figure 6C, right). Such scar lesions persisted for about 3 months (day 87) after abdominal wall resection (Figure 6D), in which fibrous scar tissue with thickened collagen fiber deposition remained in the matrix.

**Figure 6.**
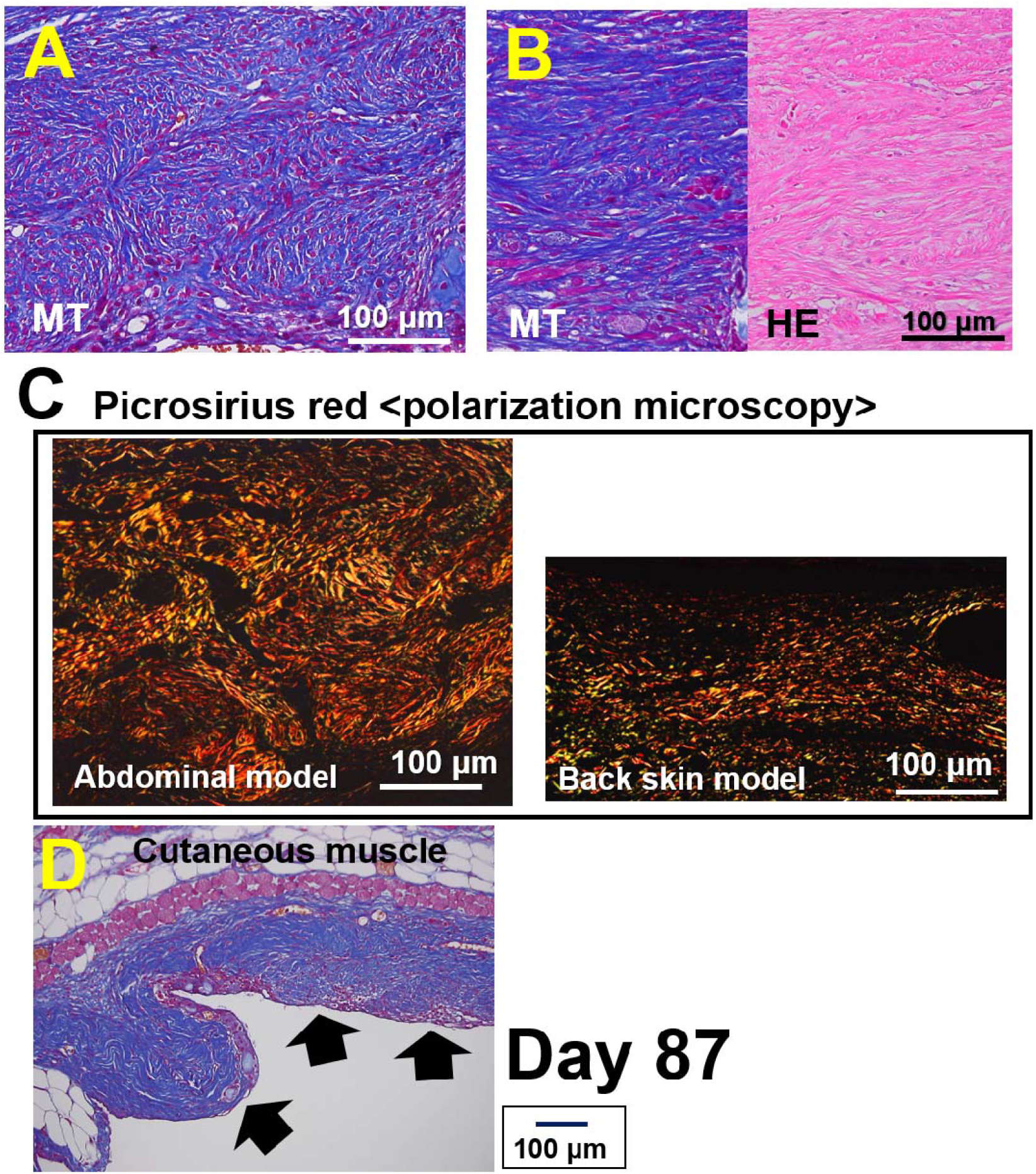
Developed and residual scar lesions in the abdominal wall model. A: Multi-nodular/spherical-like structured lesions in the abdominal wall model that developed on day 21. B: The scar lesion on day 21 shows massive collagen deposition that is hazy and hyalinized in HE stain. C: Picrosirius red staining: The fibrous lesion developed in the abdominal wall model forms a storiform pattern (left), in which thick collagen bundles are primarily stained in orange color, showing type 1 collagen. In the back skin splint model (right), thin collagen bundles were deposited in the granulation tissue on day 21, which are stained in green color, showing type 3 collagen. D: In the abdominal wall model, an apparent collagen-rich scar lesion (arrows) remained after 87 days of the study.

### Tissue weight, wound area, and lesion developments

Whether or not evaluated lesion parameters on day 21 were correlated with the initial excised tissue weight and wound area on day 0 (Figure 7). Against the tissue weight, the vertical granulation tissue length and cellular area significantly correlated (p<0.005 and p<0.01, respectively). The granulation tissue area and the collagen area tended to be related (p<0.1 in both). Against the wound area, granulation lesion length and cellular area correlated (p<0.05 and p<0.01, respectively), but no relations were with the granulation tissue area or the collagen area. The results showed that the initial wound values, including tissue weight, well reflected subsequent scar lesions in the granulation tissue on day 21.

**Figure 7.**
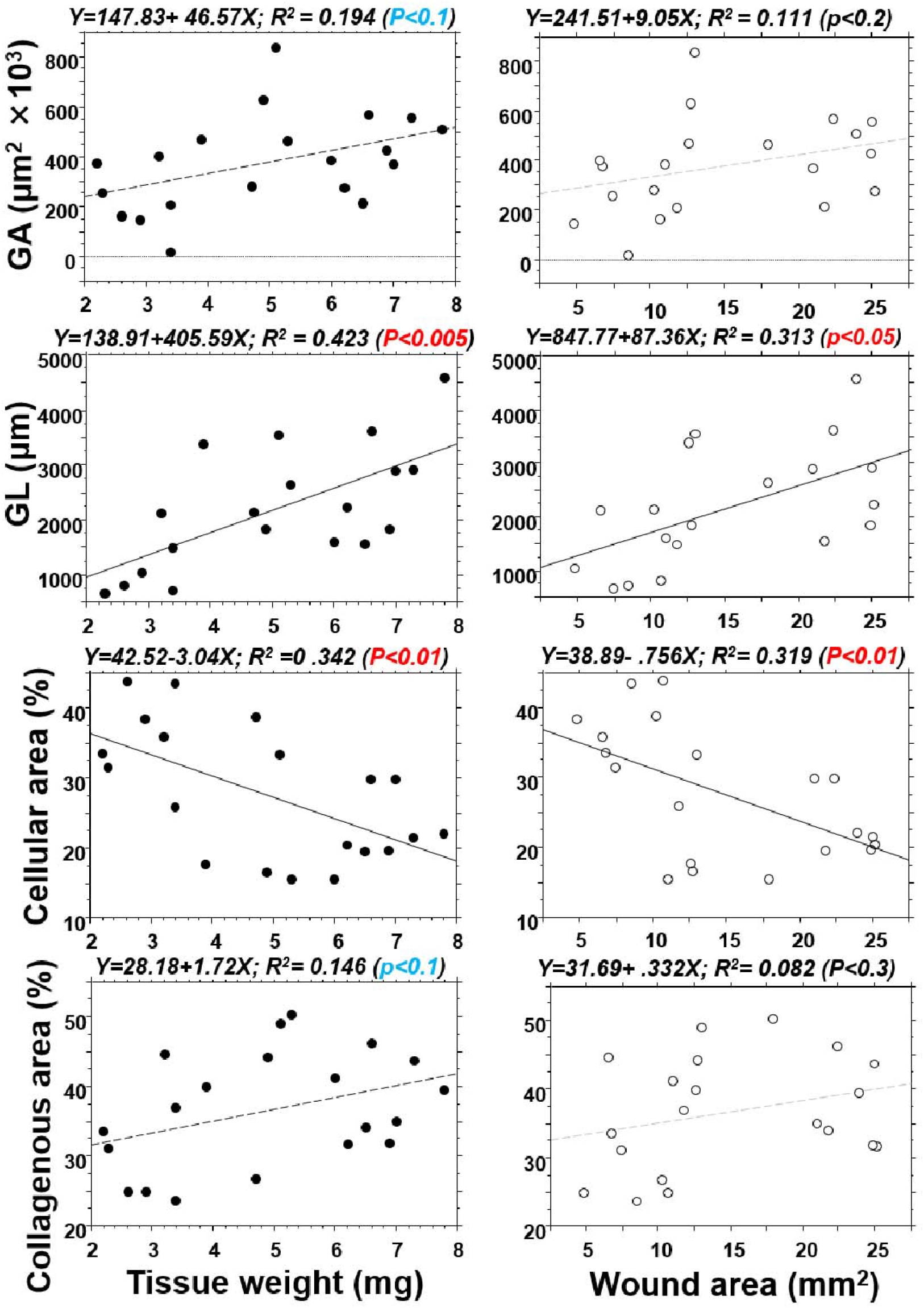
The tissue weight and wound area on day 0 and subsequent histological alterations on day 21. The correlation analysis between the original wound values (tissue weight and wound area) on day 0 and the lesion length, granulation tissue area, and collagenous area (%) on day 21. The original wound values strongly reflected subsequent scar lesions in the granulation tissue on day 21.

### How much tissue weight is necessary for achieving the developed scar lesion?

Finally, to obtain the high reproducibility for developed scar lesions in the abdominal-muscle wall model, it would be necessary to analyze the weight of the excised tissue on day 0 (Table 1). The results show that tissue weight obtained more than 5 mg may need to induce an apparent scar lesion on day 21. Therefore, more than 6 mg tissue should be excised in case of using female C57BL6/N mouse, and as a result, a more paramount and developed scar lesion may develop on day 21.

**Table 1.**
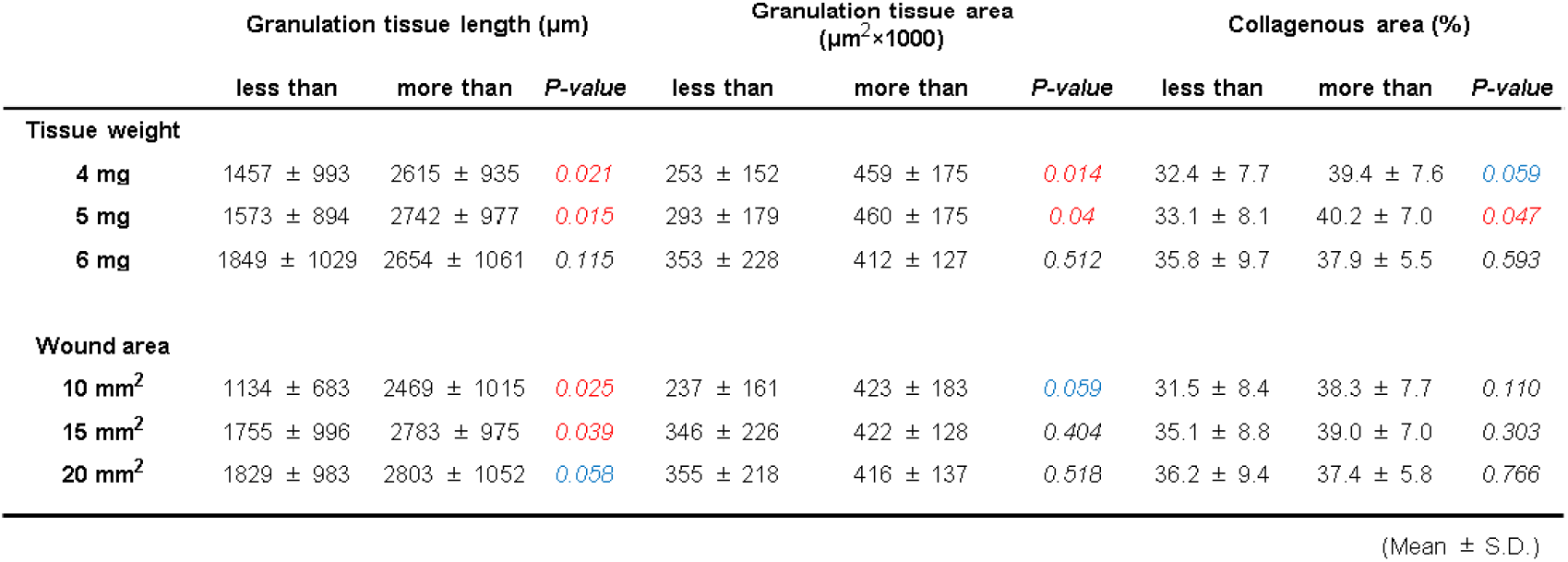
Excised tissue amounts and scar lesion formation in the abdominal model.

## Discussion

Scar lesions in humans develop not only in the skin but in other organs as well. The major pathological factors for scar lesion formation comprise a higher tensile force and chronic inflammation (Harn et al., 2019; Ogawa, 2017). However, as yet, a few models for scar tissue formation are available in rodents. Aarabi et al. (Aarabi et al., 2007) established a hypertrophic scar-forming model in mice using a tension-loading device, showing that scar tissues in the skin can develop even in mice. In other cases, fibrous granulation lesions were induced by the effect of prolonged inflammation by talc (Marchesini et al., 2020) and bleomycin (Liu et al., 2017). However, inflammation may be an additive factor for fibrosis in scaring.

In the present study, we excised a small portion of the abdominal muscle wall in mice on which skin overlayed used as a shield for the lesion against air-drying and any additional inflammation to the wounded abdominal muscle wall. No general inflammatory responses, including an increase in WBC and body weight loss, were detected in the mice after the surgery. Furthermore, NIMP-positive neutrophils and Mac-1-positive macrophages infiltrated at lower levels during the study (data not shown).

The skin wounds had healed rather quickly within 7 days after wounding. The abdominal muscle wall may be potentially/physiologically under a longitudinal static tension to some extent. Whereas, in free physiological motion after the surgery, mice twisted and stretched their body in their daily activity. Therefore, in addition to the potential muscle tension in the abdominal muscle wall, variable and complex forces could be loaded on the developing granulation tissues in the thin and soft abdominal wall that has no skeletal support.

Initially, we histologically compared the granulation tissues using the abdominal-wall excision model and the splinted back skin excision model, respectively. We had previously established the latter model (Jimi et al., 2017a). After totally excising the back skin, a round and a flexible splint was placed under the dermis, and the skin margin was fixed with it. Granulation tissue developed from the cutting edge of the fixed skin and on the facia of the back muscle. Thereby, external physical forces may load minimally on the granulation tissue developed on the wound by the splint and the back skeleton. In an early phase of granulation tissue formation on day 7, developed granulation tissue in both models revealed proliferated fibroblasts distributed in parallel. Subsequently, such lesion structure continued until day 21 of the study in the splinted skin excision model. Whereas in the abdominal-wall excision model, collagenous fibrosis progressed, and a wavy nodular structure appeared after 14 days of the study. The granulation tissues comprise many of myofibroblasts and endothelial cells. Myofibroblasts densely accumulated in the abdominal model more than that in the back-skin model. The density of the collagenous matrix increased during the 21 days of the study regardless of the model, though the value for fibrosis in the abdominal-wall excision model was about 10-times highter than in the back-skin splint model. Moreover, collagen fibers were hyalinized in the abdominal model and formed fibrous tissues consisting primarily of thickened bundles of collagen type I. These features are a signature of scar tissues (Lee et al., 2004). The described morphology and characteristics found in mice are distinct and consistent with hypertrophic scar lesions in humans (Lee et al., 2004).

In the abdominal-wall model, granulation tissue comprised neovessels, fibroblasts, and regenerative muscles in the collagen matrix. For the reproducibility of the study, we searched for a predictor for scar lesion formation using the available indexes at the time of surgery, such as the excised tissue weight and the size of penetrated wounds on the abdominal wall. Although both were significantly correlated, tissue weight may be a valuable predictive value for the scar lesions. Certainly, the correlation analysis revealed that the tissue weight was significantly interrelated with the granulation length on day 21, and tended to be related to the granulation tissue area (P<0.1), showing that tissue weight may use as a determinant of granulation tissue development after the abdominal-wall injury.

We also investigated the extent of collagen deposition in the granulation tissue leading to scar tissues using MT staining. The stained blue area highlighted was binarized and measured. The correlation analysis indicated that granulation tissue area was significantly related to the wound length and the collagen area, showing that granulation tissue size increased with fibrosis progression. Furthermore, the developed collagenous matrix in tissue was highly hyalinized, evidence for scar tissues (Lee et al., 2004). Interestingly, developed lesions showed distinct multiple nodular structures as a dominant signature for human hypertrophic scar lesions (Lee et al., 2004). Moreover, such fibrous lesions lasted for an extended period even after 87 days of the study.

The present results indicated that excised tissue weight at the time of surgery was a reliable predictive value for the granulation tissue development and consequent fibrosis on 21 days of the study. Our data indicate that when using a female C57BL6/N mouse, more than 6 mg of the abdominal muscle-wall tissue excision is necessary to get a more reliable and apparent scar lesion with advanced fibrosis.

Since the surgical procedure has been done on normal mice, active healing reactions could have occurred even though the abdominal-wall wound healing exacerbate the scaring reactions. Therefore, scar lesions developed in mice may not last as long as they do in humans. In conclusion, the abdominal-wall wound model is the first of its kind to be established for the scar lesions developed under a physiological condition without any devices. The technique is simple and reliable for studying the pathogenesis and treatment of proliferative scar lesions that develop in humans.

## Materials and Methods

### Animals

This animal study was approved by the Fukuoka University Animal Experiment Committee (No. 1812097), and study protocols complied with the institution’s animal care guidelines. C57BL/6N, female, 8-to 10-week-old mice (Japan SLC Inc., Shizuoka, Japan) were used. Basic animal treatments were followed by previously described procedures (Jimi et al., 2017a; Jimi et al., 2020a; Jimi et al., 2017b; Jimi et al., 2017c; Jimi et al., 2020b). All procedures were conducted under aseptic conditions (autoclaves, ethylene oxide gas, 70% ethanol, and povidone-iodine). Isoflurane (Wako Pure Chemical Industries, Ltd., Osaka, Japan) or pentobarbital (Somnopentyl; KYORITSU SEIYAKU, Tokyo, Japan) was used for anesthesia. Mice were monitored daily during the study. Hematological analyses, including red blood cell (RBC), and white blood cell (WBC) counts (Celltac-α, NIHON KOHDEN, Tokyo, Japan) were done in the blood drawn from the eyelids using a heparinized 75-µL capillary (Hirschmann Laborgeräte GmbH & CO., Eberstadt, Germany). At the end of the study, the mice were sacrificed by lethal pentobarbital injection and arterial hemorrhage, and wound tissues were obtained.

### Back-skin excision model (Figure 1A)

Eighteen mice were used for the back-skin excision model. To create a humanized skin wound in the mouse, our previously established internal splint method was used (Jimi et al., 2017a; Jimi et al., 2020a; Jimi et al., 2017b; Jimi et al., 2017c; Jimi et al., 2020b). In brief, under general anesthesia, a one cm circle was marked at the middle lumbar area and was excised with scissors. Then, a doughnut-shaped flexible splint was placed beneath the skin around the wound, and the splint and skin were sutured with thread (6 stitches). The wound surface was treated with 70% ethanol for 1 min. Finally, the wound was shielded with a polyurethane film dressing (Tegaderm, SUMITOMO 3M, Tokyo, Japan). A silicon vest was used to prevent thread removal by the animal.

### Abdominal wall excision model (Figure 1B)

For the abdominal wall excision model, 20 mice were used in the study for 21 days, and 10 mice were used in the study for 87 days. One day before the surgery, mice were anesthetized with pentobarbital (Somnopentyl; KYORITU SEIYAKU, Tokyo, Japan), the hair on the abdomen was removed with an electric clipper and a commercially available hair-removal cream.

On the day of surgery, mice under general anesthesia were set down in a supine position and their four limbs were fixed with pins on a corkboard. The cross point of the rectus abdominis and linea alba abdominis was marked with a dot using a marker pen. Then, a line was marked 1 cm above and 0.5 cm below the marked dot alongside the midline. Iodine was applied to the abdominal area and the skin was resected along the marked line. After separating the skin around the cut line from the abdominal muscle wall with scissors, the fascia at the umbilicus spot was pulled up and a needle with a suture was inserted. The tissue was then lifted with the thread and the spot, where the suture had been inserted, was grasped with round-top tweezers. The tissue was cut away with an ophthalmic seizer around the circular edge of the tweezers. The tissue weight was calculated.

The opened skin was placed together and sutured at upper and lower positions, which should not be attached to the wound of abdominis to avoid any side effects from foreign body reactions and to allow for determining abdominal wound healing only. The abdominal area of the wound was covered with a film dressing (Tegaderm; SUMITOMO 3M, Tokyo, Japan).

Without protecting the wounds, the mice could not survive due to their tendency to kick their abdominal organs out from their bodies. Therefore, after the surgery, a silicon vest was placed on the mice, which consists of a thin silicon sheet that was cut to mount the lower body with two holes for the lower limbs, and the upper portion of the silicon vest was stapled. The mice could move freely to drink and eat with the vest. The vests were kept on the mice for one week for the skin to be healed.

### Tissue samples

On days 0, 7, 14, 21, and 87, the mice were sacrificed under overdosed general anesthesia, and the abdominal wall, including the wound tissue, was excised and fixed in 5% buffered formalin, and paraffin blocks were prepared. The maximum appearance of cross-sectioned wounds divided into two separate blocks from one wounded tissue embed in the paraffin block was carefully determined by sectioning with the microtome. Ten serial sections cut in 4-μm thickness were initially prepared for staining with hematoxylin-eosin (HE) stain, Masson’s-trichrome (MT) stain, and immunochemical stains. To detect neovessels and myofibroblasts, rabbit anti-mouse CD31 antibody (30-time dilution: Dianova GmbH, Hamburg, Germany) and rabbit anti-mouse α-smooth muscle actin (SMA) antibody (3000-time dilution: Abcam plc, Tokyo, Japan) were used, respectively. After treatment with an enhancing reagent (EnVision Kit, DAKO Japan, Inc., Tokyo, Japan), immunohistologic localization of α-SMA was visualized by 3,3’-diaminobenzidine. Hematoxylin was used for counterstaining. For collagen distribution, paraffin sections were stained with 0.1% Sirius red in the saturated picric acid solution for one hour and examined by polarization microscopy.

### Measurement of tension in the resected abdominal muscle wall

The extent of tensile force in the abdominal muscle wall was measured. After removing the skin, the abdominal muscle was cut 1 cm in horizontal or vertical resection. After resection, the edge of the opened wound was hooked by a three-headed needle with thread. Thereafter, the tread was pulled till the wound edge returned to the equator line, for which pulling weight was measured by a tensile force measuring device. The weight was converted to the Newton unit as a tensile force on the wounds. A total of 6 mice were used.

### Morphometrical analysis

After resecting a part of the abdominal wall, penetrated wound pictures with a scale were taken, and wound areas were measure (VH-analyzer: Keyence, Osaka, Japan). Using histological pictures stained by HE and MT stains, the area of granulation tissue developed between the abdominal muscle wall and the length of granulation tissue expanded in a lateral direction were morphometrically measured with a computer-assisted software BZ-II (Keyence) (Figure 4A). To measure the area of collagenous matrix and cellular element in the granulation tissue, images of the representative lesion were taken by using 20-times objective lens in the Bio-Zero (Keyence). Collagenous matrix stained by blue and cellular elements stained by red were highlighted, binarized, and morphometrically measured by VH-analyzer (Figure 4B), and the percentage of collagen distribution and cellular element distribution in the granulation tissue was calculated.

### Statistical analysis

To compare two values, student’s t-test or Mann-Whitney U test was performed. To examine the relationship between two factors, a linear regression analysis was performed. P values < 0.05 were considered to denote statistical significance. Data are expressed as mean values ± standard deviation.

## Acknowledgments

The authors acknowledge the excellent technical support given by Fukuoka University Pharmaceutical Science students, Ms. Mina Matsuda and Mr. Masanori Fuji. We also express our gratitude for valuable discussion with Professors Drs. Hiroshi Yamada, Hiroyuki Ohjimi, and Shuuji Hara.

## Data avaiability

### Author contributions statement

S.J. was involved in tVhe animal surgery, data analysis and manuscript writing; A.S. was involved in the manuscript writing and data analysis; S.S. was involved in resources and data validation; S.K., M.M. and S.T. were involved in the data analysis and manuscript correction.

---

**Table S1.** Body weight and hematological data.

|  | Day 0 | Day 7 | Day 14 | Day 21 |
| --- | --- | --- | --- | --- |
| Dorsal skin model |  |  |  |  |
| Body weight (g) | 16.8 ± 1.1 | 17.3 ± 1.6 | 17.3 ± 1.6 | 19.0 ± 1.6 |
| WBC (x10<sup>2</sup>/μL) | 59 ± 9 | 52 ± 20 | 54 ± 13 | 50 ± 4 |
| RBC (x10<sup>4</sup>/μL) | 928 ± 21 | 938 ± 21 | 978 ± 90 | 979 ± 39 |
| Abdominal model |  |  |  |  |
| Body weight (g) | 18.7 ± 1.1 | 18.2 ± 1.3 | 20.5 ± 1.3 | 21.2 ± 1.2 |
| WBC (x10<sup>2</sup>/μL) | 40 ± 10 | 56 ± 12 | 29 ± 17 | 49 ± 12 |
| RBC (x10<sup>4</sup>/μL) | 1031 ± 23 | 900 ± 74 | 871 ± 54 | 1091 ± 94 |

Mean ± S.D.

